# Bayesian inference: The comprehensive approach to analyzing single-molecule experiments

**DOI:** 10.1101/2020.10.23.353110

**Authors:** Colin D. Kinz-Thompson, Korak Kumar Ray, Ruben L. Gonzalez

**Affiliations:** Department of Chemistry, Columbia University, New York, NY 10027 USA; Department of Chemistry, Rutgers University-Newark, Newark, NJ 07102 USA

**Keywords:** model selection, error propagation, scientific method, probability theory, kinetics, cryo-EM

## Abstract

Biophysics experiments performed at single-molecule resolution contain exceptional insight into the structural details and dynamic behavior of biological systems. However, extracting this information from the corresponding experimental data unequivocally requires applying a biophysical model. Here, we discuss how to use probability theory to apply these models to single-molecule data. Many current single-molecule data analysis methods apply parts of probability theory, sometimes unknowingly, and thus miss out on the full set of benefits provided by this self-consistent framework. The full application of probability theory involves a process called Bayesian inference that fully accounts for the uncertainties inherent to single-molecule experiments. Additionally, using Bayesian inference provides a scientifically rigorous manner to incorporate information from multiple experiments into a single analysis and to find the best biophysical model for an experiment without the risk of overfitting the data. These benefits make the Bayesian approach ideal for analyzing any type of single-molecule experiment.

## 1. INTRODUCTION

The ability to observe and characterize the biophysical properties of individual biomolecules has revolutionized the study of biological systems (38). Such single-molecule experiments avoid ensemble averaging, which removes the need to experimentally synchronize molecules and enables investigations of rare and transient molecular states. Consequently, single-molecule experiments provide unique and powerful insights into the fundamental workings of biological processes (38). Despite the mechanistically rich information contained within single-molecule data, such data are typically challenging to analyze and require extensive scientific, mathematical, and computational effort. As is the case for all scientific experiments, models play a central role in the analysis of single-molecule data. Indeed, data collected from any biophysics experiment have to ultimately be modeled according to the physico-chemical properties of the biomolecules being studied, (e.g., the molecular structure, the nature and kinetics of structural rearrangements, etc.). In the case of single-molecule biophysics experiments, this modeling process is made significantly more complex by the large uncertainties that necessarily accompany the observation of a small number of molecules for a short period of time using low signal-to-noise ratio (SNR) techniques.

Recently, methods that use probability theory and a process called ‘Bayesian inference’ have arisen as powerful tools for tackling the challenges of scientific data analysis (24), and have made a significant impact in the field of single-molecule biophysics (6). Bayesian inference formalizes the application of the scientific method to the problem of data analysis, making it an approach that naturally conforms with best scientific practices (Fig. 1) (14). Additionally, Bayesian inference-based data analysis methods require scientists to be rigorously explicit about the assumptions they make when modeling data and to fully account for uncertainties in their conclusions when the data are unclear—both important considerations when interpreting single-molecule experiments.

**Figure 1:**
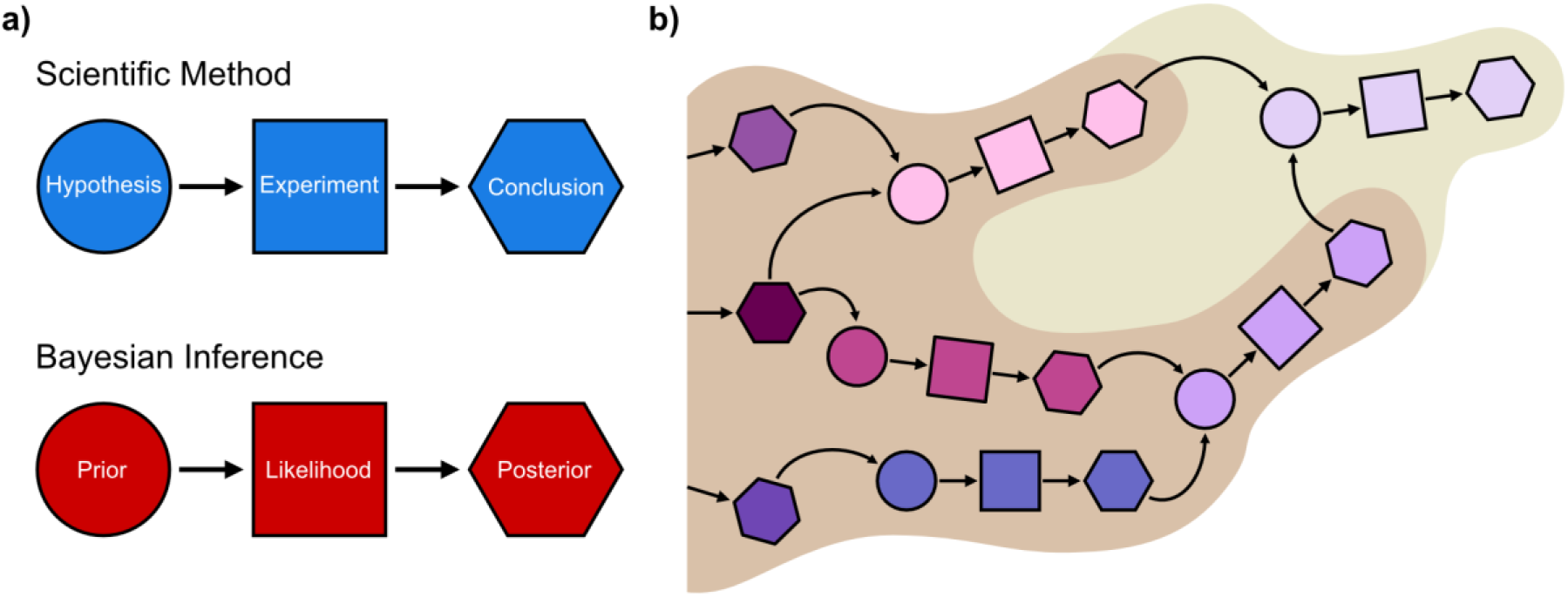
The analogy between the scientific method and Bayesian inference. **(a)** The components of a single example of the scientific method (above) show a one-to-one correspondence with those of Bayesian inference (below), revealing how the latter is just a formal extension of the former to data analysis. **(b)** The analogy is reinforced in how repeated applications of both the scientific method and Bayesian inference extend the frontier of knowledge and certainty, respectively. The area in tan shows a scientist’s knowledge (or certainty) gained by the latest application of the scientific method (or Bayesian inference), which itself is built upon previous applications.

Perhaps the most enticing and exciting aspect of Bayesian inference-based methods is the emerging possibility of using probabilities to rigorously perform ‘model selection’. When analyzing real experimental data, it is often the case that many different models are hypothetically consistent with the data. In the case of modeling single-molecule data, this problem is exacerbated by the large uncertainties inherent to the data. Bayesian inference allows one to calculate the probability that each model is the ‘best’, given both the experimental data and our previous biophysics knowledge regarding the underlying biomolecular process. Using these probabilities to select the best model and quantitatively characterize how much better it performs relative to the broader set of models under consideration is the most rigorous way to analyze an experiment.

This review addresses the question of how information can be accurately and precisely extracted from single-molecule data in a manner consistent with the principles of the scientific method. We begin by examining the role of models in the scientific investigation of a natural phenomenon. We then describe how, within the framework of probability theory, Bayesian inference uses Bayes’ theorem to extract information from experiments in a manner that is naturally consistent with the scientific method. Subsequently, we consider the specific benefits that the various terms in Bayes’ theorem (i.e., the prior, likelihood, posterior, and evidence) provide to the analysis of single-molecule experiments and provide examples of current methods that leverage these benefits. Finally, we argue for the near-future development of methods in which Bayesian inference is used to implement model selection and rigorously account for the uncertainties present in single-molecule experiments.

## 2. THE ROLE OF MODELS IN SCIENCE

The role of models in science was summarized well by John von Neumann:

> “To begin, we must emphasize a statement which I am sure you have heard before, but which must be repeated again and again. It is that the sciences do not try to explain, they hardly ever try to interpret, they mainly make models. By a model is meant a mathematical construct which, with the addition of some verbal interpretations, describes observed phenomena.” (40)

Primarily, all scientific investigations involve some combination of creating, refining, and testing these models of natural phenomena. In biophysics and related fields, for instance, one might create a structural model of a biomolecular complex or develop a mechanistic model of a biochemical reaction. The role of modeling in scientific practice is compounded when one considers that: (*i*) interpreting experimental data designed to probe such phenomena requires use of additional models to extract information that is necessary for the interpretation (e.g., models of spectroscopic signals and noise) and (*ii*) models are dependent on assumptions from associated models (e.g., structural models assume that molecules are well-modeled by point particles and bonds). Thus, to successfully model experimental data, scientists need effective models for the phenomena that they study (e.g., biophysical properties of molecules); the signals that report on, and noise that obscure, (e.g., detector signal and noise) these phenomena; and for the background knowledge on which the phenomena are conditioned (e.g., quantum mechanics) (Fig. 2).

**Figure 2:**
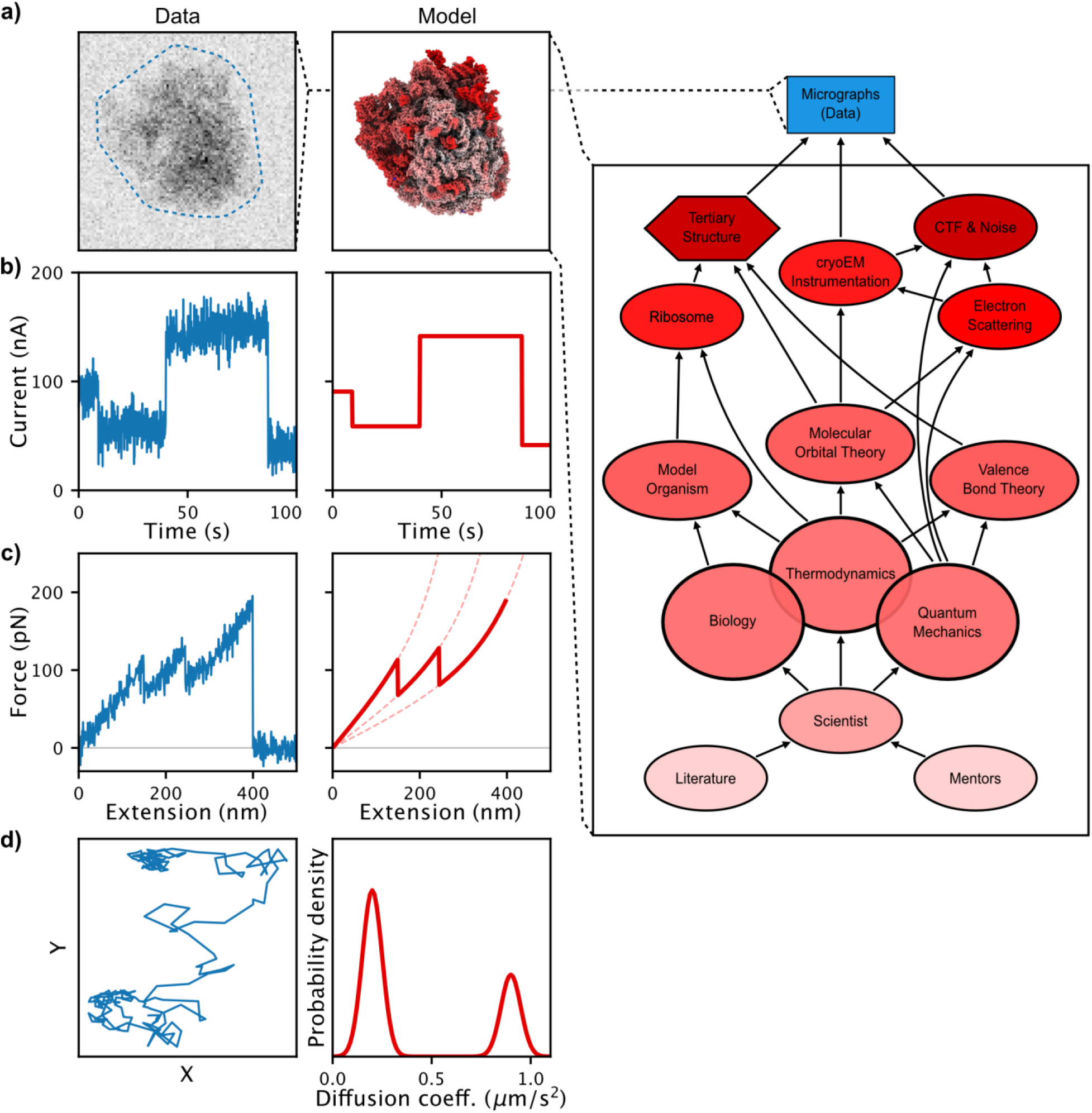
The role of models in science. Representations of simulated data (left) and a corresponding model (right) for common single-molecule studies, including **(a)** an electron micrograph probing the structure of a biomolecule (a ribosome) and the corresponding model of its structure (PDB ID: 6UZ7), **(b)** a current versus time trajectory probing the conformational dynamics of a biomolecule and the corresponding model of its conformational transitions, **(c)** a force versus extension curve probing the unfolding of a biomolecule and the corresponding model of its unfolding transitions, and **(d)** a single particle track probing the diffusion of a biomolecule and the corresponding model of the diffusion coefficient. The blowout of **(a)** shows that while scientists only aim for, and report, a portion of the model (red hexagon), the model is complex and includes noise as well as other background information (red ovals).

### 2.1 The scientific method: Experiments yield updated models

The scientific method allows us to assess which models of a natural phenomenon to trust. Models are never ‘right’ or ‘wrong’. Instead, they each provide various degrees of predictive power. Our certainty in whether a model is appropriate or not depends upon assessing that predictive power. Through this lens, a hypothesis can be thought of as a model; by performing experiments, we collect data that allows us to assess its effectiveness at explaining the natural phenomenon of interest. Based on those results, we can ‘update’ the model to better represent the phenomenon in the future, or move on to a different model.

Another way to think of the scientific method is to consider two separate models, say *M*_1_ and *M*_2_, that are the same except for the value of a single parameter. For example, *M*_1_ and *M*_2_ might represent slightly different conformations of a biomolecule, and given our prior knowledge of this biomolecule, our initial model of its conformation might be that both *M*_1_ and *M*_2_ are equally reasonable. By performing an experiment, we might determine that *M*_1_ is better able to describe the observed data than is *M*_2_. Thus, performing the experiment can be thought to have updated our conformational model to favor *M*_1_, which has more predictive power. By extrapolating this process to models that differ by many parameters or that are conceptually distinct, it becomes apparent how implementing the scientific method generally enables experiments to yield updated models of natural phenomena (Fig. 1).

### 2.2 Models in single-molecule studies

Although single-molecule experiments are incredibly rich sources of data, observing and trying to characterize the behavior of a set of individual molecules complicates the modeling process. This is because, rather than using a single model to describe the average molecular behavior as is done in an ensemble experiment, the behavior of every individual molecule in a single-molecule experiment must be separately modeled and then those individual models must somehow be integrated into a collective model that describes the overall behavior of the biophysical system.

Moreover, uncertainties originating from the sample, the instrumentation used to collect the data, and the analysis of the collected data further compound the challenges associated with modeling single-molecule data. Compositional and spatial heterogeneities in the sample, such as differences in post-translational modifications and in the local molecular environment, respectively, can make any one observed biomolecule different from the other observed biomolecules. Furthermore, samples using reporter molecules, such as fluorophores in single-molecule fluorescence experiments, can exhibit heterogeneous signaling dynamics (e.g., photophysical effects such as photoblinking or photobleaching). The presence of these heterogeneities across the individual molecules complicate data modeling, and, thus, the analysis process. Additionally, the SNR of data from an individual molecule is generally low, despite the high sensitivity of the instruments used to collect these data. Such low SNR makes it difficult to model data with a high degree of confidence. These instruments also generally have limited observation times and/or throughputs, both of which make it difficult to collect a statistically relevant amount of data. A further complication is that it remains theoretically unclear whether the data from a single molecule observed over a long period of time are equivalent to the data from multiple individual molecules observed over a shorter period of time (i.e., whether biological systems are ‘ergodic’), an assumption that is implicit in many data analysis methods. Finally, the models used to appropriately describe the behavior of individual molecules are often not well-developed, and are themselves a subject of active research (19).

Regardless of these complications, the data recorded from the individual molecules in a single-molecule experiment have to be modeled in order for conclusions to be drawn about the biomolecular process under investigation. Fortunately, the scientist often has prior knowledge of the biomolecular system that can inform their modeling. For example, knowledge of the primary and/or secondary structure of a molecule can inform tertiary structural modeling. In the following section, we will show how to use probability theory to apply a model and extract the relevant information in a mathematically rigorous manner that accounts for all of the uncertainties in the modeling process while making use of such prior knowledge.

## 3. USING PROBABILITY THEORY TO MODEL EXPERIMENTS

In 1946, Cox showed how a ‘probability’, *P*, can be understood as an extension of formal logic that quantifies the certainty in a scientific statement (5, 14). A statement has *P* = 0 if false and *P* = 1 if true. A fractional value of *P* between zero and one corresponds to the certainty that the statement is true. For example, consider the model defined by the statement “every molecule in the ensemble is in the same conformational state” (*M*_*same*_). Even before performing an experiment to test it, we know that *M*_*same*_ corresponds to a system with extremely low entropy, and, according to the second law of thermodynamics, it is very unlikely that *M*_*same*_ is true. Probability theory allows us to write this as *P*(*M*_*same*_) ≈ 0. Of course, this assessment is based on more than a century of biophysics knowledge; so, to be transparent about the scientific knowledge incorporated into our certainty in that statement, we must write that it conditionally depends upon the model of our biophysics knowledge (*M*_*biophysics*_). This conditional probability should thus be explicitly written as *P*(*M*_*same*_|*M*_*biophysics*_) ≈ 0, where the vertical bar reads as the word ‘given’.

Although they are often not explicitly acknowledged, all scientific statements are conditionally dependent upon the scientist’s notions of background models. For instance, it is generally true that all biophysicists’ analyses adhere to the laws of thermodynamics; so, there is little need to explicitly acknowledge that dependence in an analysis. Similarly, it is generally true that all probabilities have conditional dependencies, but when those dependencies are obvious or seem unimportant there is little need to explicitly acknowledge them. Regardless of whether such conditional dependencies are explicitly acknowledged in an analysis or not, every analysis is still dependent upon them. However, to some, acknowledging the conditional dependence of an analysis upon, for example, one specific scientist’s *M*_*biophysics*_ is seen as incorporating a subjective, non-scientific element into an analysis. It is important to note, however, that this merely reflects a more general, though unfounded criticism of the role of conditional dependencies in the scientific method itself. Fortunately, while two scientists may have learned biophysics from different sources, and thus technically have different *M*_*biophysics*_s, the collective body of knowledge that defines a field like biophysics compels two scientists with an equivalent exposure to the field to have effectively equivalent *M*_*biophysics*_s. The proof of this is that two well-informed scientists endeavoring to perform the same experiment to test the same model undoubtedly reach the same conclusions to a high enough precision that science is reproducible.

The most beneficial aspect of the correspondence between probabilities and scientific statements is that probability theory can be used to quantify the effects of an experiment on our certainty in a scientific statement. For example, consider a model, *M*_*conformations*_, that attempts to quantify the conformation of each biomolecule in a homogeneous ensemble. The set of parameters of *M*_*conformations*_, {θ}, might be the Cartesian coordinates of all the atoms in all of the biomolecules in the ensemble. Given our *M*_*biophysics*_, we may have some idea before performing an experiment about the particular values of {θ} that are reasonable (e.g., atoms are not closer to each other than 1 Å). Thus, in the context of *M*_*conformations*_, the probability that the ensemble of biomolecules exists in one particular set of conformations is *P*({θ}|*M*_*conformations*_, *M*_*biophysics*_). Because *P*({θ}|*M*_*conformations*_, *M*_*biophysics*_) can be formulated before an experiment is performed, it is called a ‘prior probability’. After performing an experiment that is designed to probe the conformations of the biomolecules (e.g., measuring fluorescence resonance energy transfer (FRET) efficiencies (E_FRET_s) with a single-molecule FRET (smFRET) experiment), the set of data, {D}, that was collected and processed using a model of the experiment, *Mexperiment*, will update the prior probability of a particular {θ} to a ‘posterior probability’ value, which is written *P*({θ}|{*D*}, *M*_*experiment*_, *M*_*conformations*_, *M*_*biophysics*_). This posterior probability is also the probability of a particular {θ}, but is conditionally dependent upon the newly observed, experimental data obtained and processed according to *M*_*experiment*_ (e.g., observed E_FRET_ values are assigned to particular conformational states using a separate experiment, such as a cryogenic electron microscopy (cryo-EM) study). In the following section, we will discuss exactly how {D} is used to update a prior probability into a posterior probability.

### 3.1 Bayesian inference: Applying the scientific method to data analysis

Bayesian inference is the process of using probability theory to model and analyze experimental data. Specifically, it is the application of Bayes’ theorem to estimate the posterior probability for the values of a set of model parameters, {θ}, conditionally dependent on experimental data, {D}, for a particular scientific model, *M*. The goal of any analysis is to find the ‘best’ {θ} for the *M* to describe the natural phenomenon, and to judge this using the observed {D}. Practically, it is often the case that many different {θ}’s will yield a reasonable version of *M*, and this is especially true if {D} contains significant statistical uncertainty. The solution is to use Bayes’ theorem to calculate the posterior probability, *P*({θ}|{D}, M), of every possible instance of {θ}

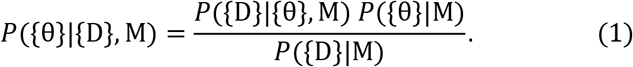

As explained above, obtaining the posterior probability distribution, sometimes simply called the posterior, is the goal of this inference process. In the numerator, the term *P*({*D*}|{θ}, *M*) is called the likelihood function, or simply the likelihood, and *P*({θ}|*M*) is the prior probability distribution, or simply the prior. In the denominator, *P*({*D*}|*M*) is called the evidence function, or simply the evidence. The evidence may be rewritten as

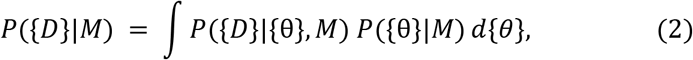

where the integral is taken over all possible values of the set of {θ}. This type of integration is called ‘marginalization’, because it removes the dependence on {θ}. Thus, the evidence can be interpreted as the probability of observing the {*D*} regardless of the exact values of {θ} for the *M*; because of this interpretation, the evidence is sometimes called the ‘marginal likelihood’. This means that Eqn. 1 involves only the prior and the likelihood, and that Bayesian inference is performed by choosing the *M* (which involves defining the prior) and then collecting the {*D*}—after which the resulting posterior yields insight into the phenomenon being modeled by *M*.

One of the most powerful aspects of using Bayesian inference to analyze an experiment is that it is analogous to using the scientific method (c.f., Section 2 and Fig. 1). Forming a hypothesis to test with the scientific method is equivalent to choosing a model and defining the prior for Bayesian inference. Analyzing the results of an experiment to reach an updated conclusion about the hypothesis is equivalent to using the likelihood to obtain the posterior. In this sense, Bayesian inference allows scientists to rigorously extend the scientific method into the realm of analyzing their data, and vice versa. In addition, the scientific method relies upon multiple, interconnected models to investigate a natural phenomenon (c.f., Section 3) and Bayesian inference explicitly details how the analysis of an experiment depends on those models. This mirroring of the scientific method is what makes Bayesian inference such powerful analytical tool. In the following sections, we will discuss the various terms in Bayes’ theorem, the contributions they make to Bayesian inference, and the distinct benefits they provide to the analysis of single-molecule experiments. In each section, we have highlighted specific examples of analytical tools, algorithms, and/or software packages in which the term described in that particular section has been used to great effect in the analysis of single-molecule data. Given that we are only able to highlight a limited number of specific examples, we point the interested reader to additional specific examples in the Related Resources section at the end of this article.

### 3.2 The prior: Quantifying the hypothesis

Before performing an experiment, a scientist generally has prior knowledge about the natural phenomenon under investigation that led them to develop the hypothesis they are testing. The prior, *P*({θ}|*M*), quantifies this knowledge about {θ} for the *M* being tested. For instance, when trying to determine the rate constant for a biomolecular conformational change, a very reasonable prior based on our *M*_*biophysics*_ would be to stipulate that the rate constant has a non-zero probability for being in the range between 3.2×10^−8^ s^−1^ (less than a year) and 10^14^ s^−1^ (more than a bond stretching time) and a probability of zero for being outside that range. In practice, our prior knowledge about a particular biomolecular system often allows us to specify priors with more information than the very loose range in this example.

The use of a prior provides many benefits to the analysis of single-molecule experiments, but two of them are particularly powerful. First, the use of priors enables precise analysis of very small amounts of data. This is because any amount of collected data, even a single data point, will update the prior into the posterior. This is extremely advantageous for the analysis of single-molecule experiments, which frequently yield relatively small datasets. Second, priors provide a coherent, mathematical framework for incorporating information from previous experiments into the current analysis—even if those experiments were performed with different experimental techniques (e.g., refinement of a cryo-EM structure using structural homology (11, 20)).

Just as formulating a sound hypothesis is the art of the scientific method, choosing an appropriate prior is the art of Bayesian inference. For example, priors that describe years of knowledge about the signals, noise, and transition kinetics that are typical of the E_FRET_ versus time trajectories (E_FRET_ trajectories) recorded in smFRET experiments are used in the Bayesian inference-based smFRET analysis methods vbFRET, VB-HMM-TS-FRET, ebFRET, bl-ICON and hFRET (2, 13, 29, 34, 39). Notably, the use of a prior for the transition kinetics in these methods ensures that a posterior quantifying the transition kinetics exists, even if no transitions occur in the E_FRET_ trajectory being analyzed. Essentially, the absence of any observed transitions is able to provide an upper-limit for the transition rate; non-Bayesian inference-based methods cannot reach this conclusion.

Similarly, in the Bayesian inference-based cryo-EM single-particle analysis (SPA) method RELION, priors are used for the Fourier components (*i.e.*, the coefficients of the spatial frequencies) in the density map reconstructed from electron micrograph images (33). By using Gaussian distributions centered at zero for these Fourier components, the use of priors in RELION enables high-resolution mapping of the electrostatic potential of a molecule while simultaneously avoiding spurious noise from the high spatial frequencies where there is little structural information present in the raw data. Work to incorporate more information into these priors is underway, for instance by including information about the inherent spatial frequencies found in all biomolecular structures (17).

Once a prior is specified, an experiment can be thought of as acting via the likelihood to redistribute the probability of {θ} specified by the prior to where it is most consistent with the collected data; this new, updated distribution is the posterior. Importantly, the amount of data collected in an experiment typically overwhelms the information content in the prior, and dominates the posterior result; otherwise, it would be unclear why the scientist thought the particular experiment should have been performed in the first place (an idea explored in Bayesian experimental design (3), but beyond the scope of this review). Moreover, the incorporation of incorrect knowledge or ‘bad’ information into the prior does not pose a major concern, because, beyond the transparency requirement of specifying the actual background information used in the analysis as a conditional probability, Bayesian inference-based model selection should be used to simultaneously test multiple models (c.f., Section 4). Such an approach should quickly eliminate models with bad prior choices, and yield the best description of the natural phenomenon being investigated.

An occasional criticism of Bayesian inference, and particularly priors, is that it can introduce a ‘non-scientific bias’ into an otherwise ‘objective’ analysis. The use of priors does not introduce bias into a scientific model, however, it is instead part of the mathematical statement of the ‘biases’ that already exist in the scientific investigation (c.f., Section 3); all analysis methods, Bayesian or not, include such ‘biases’. In fact, it is possible to employ priors in a Bayesian method that express the biases inherent to non-Bayesian methods, such as maximum likelihood estimation (MLE) method (c.f., Section 3.3). By ignoring the existence of the prior, as well as the posterior and evidence, such non-Bayesian methods do not fully enjoy the benefits of probability theory, including the abilities to mathematically adhere to the tenets of the scientific method (Section 3.1), properly quantify the uncertainty in the model of the experiment (Section 3.4), and perform model selection to determine the ‘best’ model and avoid overfitting (Section 4).

### 3.3 The likelihood: How an experiment relates to a model

The likelihood function, *P*({*D*}|{θ}, *M*), can be thought of as the mathematical equivalent of the experiment used to test *M* (Fig. 1). Assuming that *M* is the ‘true’ representation of the natural phenomenon being studied, the likelihood is the probability of observing a particular {*D*} in the experiment, given that {θ} comprises the ‘true’ parameters of *M*. For the analysis of single-molecule experiments, it can be quite challenging to devise and write down the likelihood function, because its mathematical form must encapsulate the model itself. Identifying and deriving suitable likelihoods that capture the complex and/or heterogeneous behavior of an individual molecule for the many different experimental single-molecule techniques is often the limiting factor in the Bayesian inference-based analysis of single-molecule experiments, and is often itself the subject of intense theoretical study (10). For instance, the method BIASD is used to analyze time series data from single-molecule experiments where the underlying molecular dynamics are faster than the instrumental time resolution (18); hFRET is used to analyze time series data from single-molecule experiments where the molecules exhibit heterogenous kinetics (13); and bioEM is used to analyze structural data from cryo-EM experiments where the molecules exhibit heterogeneous conformations (4). All of these examples of Bayesian inference-based methods use specialized likelihood functions for overcoming the complexities present in single-molecule data.

It should be noted that the likelihood is also used extensively in non-Bayesian inference-based data analysis methods—particularly in MLE-based methods. In MLE-based methods, the {θ} that yields the highest value of the likelihood for the observed {*D*} is used as a point estimate of the model of the underlying phenomenon. Because MLE does not acknowledge the uncertainty in {θ}, MLE-based methods suffer from severe overfitting (p. 434 in 1) and can be inappropriate for analyzing data from single-molecule experiments where the uncertainties can be quite large. Additionally, while the likelihood function is the conditional probability of {D} based on a particular {θ}, the objective of modeling a natural phenomenon according to the scientific method is to determine the optimal {θ} based on {D}. Thus, MLE-based data analysis methods address the reverse problem to what the scientific method aims to solve. It is worth noting that, with a prior that is independent of {θ} (i.e., a ‘flat’ prior), the posterior is proportional to the likelihood. In this case, the maximum of the posterior, which can be found using the Bayesian technique called maximum a posteriori (MAP) estimation, is numerically the same value as the point-estimate found with MLE. Nonetheless, non-Bayesian methods such as MLE miss out on all the benefits that using Bayesian inference provides for single-molecule data analysis (c.f., Sections 3.2–4).

### 3.4 The posterior: Updating the model after performing the experiment

The posterior, *P*({θ}|{*D*}), can be thought of as the quantification of how the experimental {*D*} updates our certainty of the initial hypothesis (i.e., the prior) in terms of the model parameters in {θ}. In essence, all data analysis methods that are consistent with the scientific method strive to obtain the posterior—regardless of whether they acknowledge it or not. While some analysis approaches simply attempt to estimate the single ‘best’ {θ} to explain the experimental data (e.g., MAP), the posterior provides the probability for all possible values of {θ}. As such, this makes the reporting of the entire posterior tedious or, if it has no analytical form, impossible. Thus, common approaches to reporting posteriors include providing the credible interval, which describes the range of {θ} that contains a certain percentage (e.g., 95%) of the posterior probability. It can also be useful to provide summary statistics of the posterior, such as expectation values and variances of {θ} from the posterior.

Despite the benefits it provides to the analysis of single-molecule experiments, fully implementing Bayesian inference has historically been quite difficult in practice. Specifically, this is because of the mathematical challenge of deriving analytical equations for the posterior and the computational cost of evaluating numerical solutions for posteriors without analytical solutions (1, 24). There are several approaches that directly address these challenges. One approach is to only consider models that yield analytical solutions. But this approach limits the variety of priors and likelihood functions that can be used, which may limit the ability to represent the actual scientific knowledge used to create the model. Instead, given modern computational resources, the more appropriate approach of numerically calculating the posterior is now easily achievable. The standard approach is to use a Markov chain Monte Carlo (MCMC) sampling variant (8, 9, 12, 25), which will yield the full posterior and are exact to an arbitrary precision that depends on the amount of sampling (1).

Another general and computationally feasible approach is to use methods that yield tractable approximations of the posterior. Of these, the standard is the ‘Laplace approximation,’ where the posterior is assumed to be a multivariate Gaussian distribution centered at the maximum of the posterior (*i.e.*, the MAP point) with a variance calculated from the curvature of the posterior at that point (1)—a very reasonable approximation as a consequence of the central limit theorem when there is enough data in {*D*}. Importantly, the Laplace approximation is not much more computationally intensive than finding the MAP point, but still provides a full, although approximate, posterior. This suggests that using flat priors, finding the MAP point, and then calculating the Laplace approximation of the posterior will easily allow any MLE-based method to be converted into a Bayesian inference-based method. Thus, the Laplace approximation allows both current MLE- and MAP-based methods to be easily extended to obtain an approximate posterior, and, consequently, the evidence (c.f., Section 3.5).

In cases where the Laplace approximation provides a poor approximation of the posterior (e.g., single-molecule experiments with a small number of datapoints) more mathematically rigorous approximation methods, such as a variational approximation, can be used. The variational approximation used in variational Bayesian (VB) inference is the same as that used in quantum mechanics. This approach depends on the fact that any approximation of the true posterior will have an evidence value that is a lower bound for the true evidence value (i.e., the evidence lower bound, or ELBO), achieving equality when the approximate posterior is equivalent to the true posterior (1). Thus, in VB inference, the best approximations of the true posterior are found by searching for the maximum value of the ELBO. The first use of VB inference in single-molecule biophysics was with vbFRET, which uses VB inference to yield a tractable, analytical form of the posterior for a hidden Markov model (HMM) in order to model the E_FRET_ trajectories recorded in smFRET experiments (2). Because the VB inference approach is both accurate and efficient, it has found widespread use in many single-molecule biophysics methods such as ebFRET (39), hFRET (13), vbSPT (30), VB-HMM-TS-FRET (29) and others (15, 36). In addition to these benefits, the real power of VB inference methods is that they also provide an estimate of the true evidence (in the form of the ELBO). Thus, they can be used to perform model selection (Section 4).

### 3.5 The evidence: Evaluating the effectiveness of a model

The evidence, *P*({*D*}|*M*), is perhaps both the most powerful and overlooked term in Bayesian inference. It provides the probability that the observed {D} could have come from the *M* being tested, regardless of the specifics concerning {θ}. As discussed in Section 3.1, the evidence is obtained by marginalizing out every possible value of {θ}, and thus can be thought of as being agnostic towards their ‘true’ value. Interestingly, because this marginalization is an integration performed over all of the model parameters, the more parameters included in a model, the more the predictive power of the model is diminished. The intuition behind this mathematical phenomenon comes from the fact that, while a model with a large number of parameters may describe the particular observed dataset very well, it is also flexible enough to account for a large number of other possible datasets. In this sense, the overall probability that the observed dataset originated from the model in question (*i.e.*, the evidence) is diluted by the existence of the large number of plausible datasets that could have been generated by the model (1). Thus, the evidence protects against overfitting by balancing the ability of a model to explain the observed data and its ability to generate only the observed data, thereby favoring models with the highest predictive-power and simultaneously the fewest parameters. This is a very attractive property for scientists, as it is mathematically equivalent to Occam’s razor, which states that the most parsimonious model is the ‘best’ model.

Unfortunately, just as with the posterior (see Section 3.4), the evidence is often difficult or impossible to directly calculate. As such, it is often ignored in many data analysis methods. For instance, it is unimportant when finding the MAP solution of a posterior (*i.e.*, the point estimate of the location of the maximum of the posterior), because the value of the evidence is independent of {θ} and thus will not change the location of the maximum. Nonetheless, just as with the posterior, there are a number of methods available for approximating the evidence. When using the Laplace approximation (see Section 3.4), for example, the evidence has the analytical form corresponding to a multivariate Gaussian posterior (1). Similarly, when using VB inference (see Section 3.4), the ELBO provides a measure of the true evidence. In particular, if great care is taken to find the best possible variational approximation of the posterior, the ELBO will achieve the true value of the evidence to within arbitrary precision. Thus, in contrast to an approximation of the evidence that, by construction, will never be correct, if properly treated, the ELBO can be used as an estimate of the true value of the evidence.

A further, very rough approximation of the evidence is the Bayesian information criterion (BIC), which approximates the evidence of the Laplace approximation in the asymptotic limit that there are so many data points that both the prior and the variance of the posterior can just be ignored. Considering the relatively limited number of datapoints in single-molecule experiments and the correlations present in {θ}, the assumptions that lead to the BIC (or the conceptually similar, but ad hoc, Akaike information criterion (AIC)) should not be used for single-molecule data analysis (2). The full Laplace approximation out-performs the BIC in model selection, captures the correlations in {θ}, and only requires minimal additional computation beyond the MAP solution (1, 26). It is worth noting, however, that rather than obtaining an approximation of the evidence, it is possible, albeit computationally expensive, to numerically calculate the exact value of the evidence using MCMC sampling with a method called thermodynamic integration (21).

Because of the large uncertainties associated with single-molecule experiments, the evidence is a particularly powerful tool for analyzing experiments performed at single-molecule resolution. By marginalizing out all of the possible {θ} from *M*, the evidence quantifies how consistent the observed single-molecule data is with *M*. For instance, vbFRET (and other similar methods (13, 29)) models an E_FRET_ trajectory collected in an smFRET experiment with a series of HMMs employing an increasing number of hidden states (2). Of these, the HMM with the largest ELBO corresponds to the most parsimonious model appropriate for the observed E_FRET_ trajectory, and is taken to be ‘best’ model for that E_FRET_ trajectory. By using the evidence, HMMs with more hidden states than are required to explain the data are ‘penalized’, which allows this ‘maximum evidence’ approach to avoid overfitting (Fig 3). Similarly, the maximum evidence approach is routinely used in single particle tracking (SPT) experiments to choose between competing models of diffusion based on particle trajectories of limited length (27, 28, 31, 35, 37).

**Figure 3:**
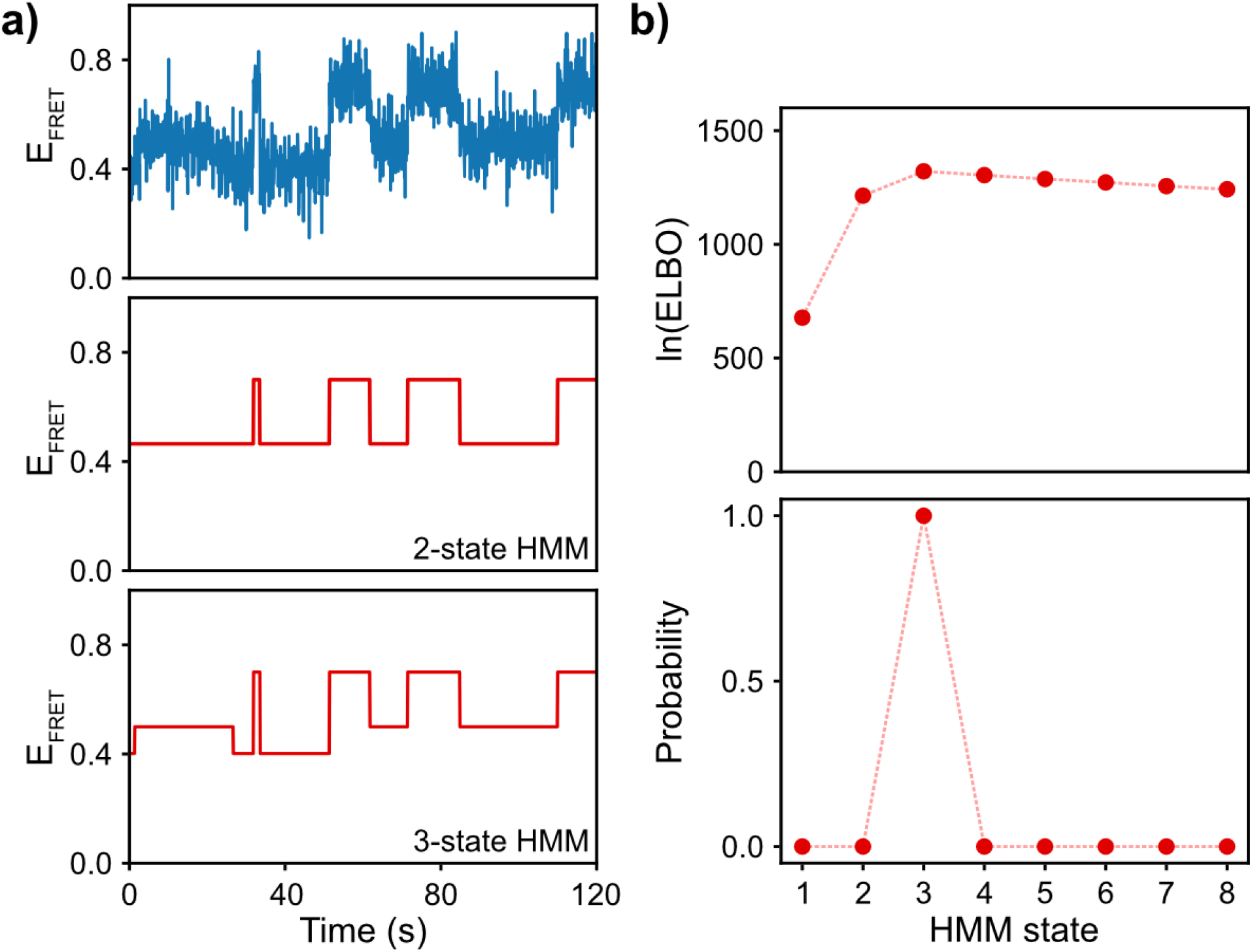
Bayesian model selection. **(a)** Representation of a typical E_FRET_ trajectory (top) and the corresponding 2-state (middle) and 3-state (bottom) HMMs for the trajectory, as analyzed by vbFRET. **(b)** The log of the ELBOs for HMMs with increasing number of states (as calculated by vbFRET) shows a peak at the 3-state model (above), and decays slowly as more states are added. Upon using these ELBOs to calculate the posterior probability for these models (below), it is clear that the 3-state model is overwhelmingly more probable than the others.

When modeling any biophysical process, it is worth mentioning that the models developed do not account for every possible experimental complication found in the data. In such cases, one often finds that the ‘maximum evidence’ approach is difficult to implement. For instance, there might be several models with evidences that are probabilistically too close to each other to choose any of them as having the ‘largest’ evidence. In Section 4, we discuss how to determine the ‘best’ model indicated by the evidence using probability theory to account for the uncertainty present in the data and models.

## 4. MODEL SELECTION: DETERMINING THE BEST MODEL USING PROBABILITY THEORY

The goal of the scientific method is to perform experiments in order to test a hypothesis that, on some level, will ultimately inform upon more than the experiment itself. For instance, an understanding of the role that the conformational dynamics of a biomolecule plays in a particular biochemical reaction informs more broadly upon biomolecular function in general. Practically, this means that, at some point during an investigation, a decision must be made about what the ‘best’ model for the phenomenon being studied should be in order to inform upon other phenomena. In Section 3.5, we discussed how the evidence, *P*({*D*}|*M*), quantifies the predictive power of a model, and showed how Bayesian-inference based methods, such as vbFRET (2) and others (13, 15, 27–31, 35–37), can utilize the maximum evidence approach to choose the ‘best’ model for the data. The maximum evidence approach, however, fails to account for the uncertainty from the limited amount of data collected during an experiment. For instance, how does one select, as is often the case in single-molecule experiments, between models with effectively the same evidence value? The answer is to take the ideas developed in the sections above one step further and make this determination in a manner consistent with probability theory by again using Bayesian inference.

Bayesian model selection (BMS) essentially entails performing a second round of Bayesian inference where the evidences for each model are used as likelihoods to calculate a posterior probability for the models themselves (1). In practice, a scientist can assign a model prior probability to each model under consideration that it is the ‘true’ model as *P*(*M*_*i*_|*M*_*biophysics*_), where *M*_*i*_ is the *i*^th^ model under consideration, such that ∑_*i*_ *P*(*M*_*i*_|*M*_*biophysics*_) = 1. If there is no reason to favor any *M*_*i*_ over the others, then these model priors should all be equal; models not considered or not imagined, given a scientist’s *M*_*biophysics*_, can be thought of as having a model prior probability of zero. Using the evidences for each model, *P*({*D*}|*M*_*i*_, *M*_*biophysics*_), the model posterior probability for *M*_*i*_, *P*(*M*_*i*_|{*D*}, *M*_*biophysics*_), can then be calculated as

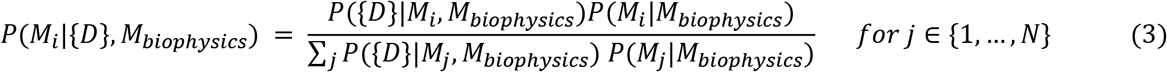

Comparing Eqn. 3 to Eqn. 1 demonstrates that BMS is a form of Bayesian inference, and so all of the benefits of using priors, likelihoods, and posteriors detailed in Section 3 also apply here.

The model posterior is the object of BMS, as it eliminates the difficulties of trying to arbitrarily assess whether the evidences for two models are effectively the same or not; this also addresses the issue of ‘plateauing’ evidences often found in maximum evidence methods such as vbFRET (2). There are two approaches to deciding which model to use after performing model selection and calculating the model posterior. First, the model with the largest model posterior value can be chosen. Second, a probability threshold can be used (and set before performing the experiment) where the scientist can decide that the experiment was ambiguous if none of the models surpass the threshold (e.g., greater than 0.95). If no model surpassed the threshold, a subsequent experiment would have to be performed, perhaps with a different technique to provide distinct information or with more data to be collected, in order to distinguish between the models. Thus, the uncertainty quantified with the posterior in BMS allows the scientist to assess the effectiveness of the experiment and subsequent analysis.

We believe single-molecule experiments are best analyzed in this manner, because the extensive use of probability theory enables scientists to easily deal with the statistical uncertainty and other related problems faced in single-molecule experiments (*c.f.*, Section 2.2) with a unified and comprehensive framework. Single-molecule analysis methods that currently employ evidences (or ELBOs, for VB approaches) can be easily extended to perform BMS by using those evidences with Eqn. 3 (Fig 3). Thus, BMS can be used to extend current methods to determine the number of hidden states in an smFRET study, the best structural model for each conformational class in a cryo-EM study, the best model for diffusion dynamics, etc. (c.f., Section 3.5). Currently, BMS is used to determine the presence of a change-point in a signal-vs-time trajectory (7), the best forcefield to be use in the construction of structural models (10), and even whether a noisy fluorescent image corresponds to the molecule of interest or is ‘junk’ (32). The list of models that can be imagined to analyze single-molecule experiments is nearly endless, and, thus, so too is the number of applications for BMS in single-molecule biophysics.

## 5. CONCLUSION

It is clear that implementing Bayesian inference, even approximately, is extremely powerful for single-molecule data analysis, and has enabled deep insight into biomolecular systems through the rational and judicious use of priors, likelihoods, posteriors, and evidences. Not only is Bayesian inference incredibly effective as an analysis tool for single-molecule experiments, but it is also the most optimal tool, as it enables a scientist to account for the large uncertainty in single-molecule data. Additionally, it allows a scientist to do so in a way that is rigorously consistent with the scientific method, to be transparent about the underlying assumptions used in the modeling, and, most importantly, to select the best model of a phenomenon in a quantitative, scientific manner. It is worth noting that, while we have focused our attention on the analysis of single-molecule biophysics experiments, data from experiments in practically all scientific fields exhibit a finite signal-to-noise and are composed of a finite number of data points (c.f., Section 2.2). The universal applicability of the Bayesian approach to analyzing data therefore stands to benefit scientific exploration in virtually all fields.

Unfortunately, despite the great advantages that they offer, many of the Bayesian inference-based methods described above have not yet been widely adopted. While this may be at least partly due to the misconception that Bayesian inference, and particularly the use of priors, might introduce ‘non-scientific bias’ into an analysis (see Section 3.2), it is clear that further work is still required to make Bayesian inference-based methods more accessible, computationally efficient, and capable of modeling more complex single-molecule data. Fortunately, recent progress in the field demonstrates that addressing these shortcomings is a very active area of research (13, 16, 22, 23, 41). Implementing the BMS approach as we have described in Section 4 makes use of all the benefits that probability theory affords and is an exciting avenue to explore for single-molecule analysis methods under current or future development. It is our hope that this review will encourage others to use currently available Bayesian inference-based methods in their single-molecule data analysis pipelines and inspire them to develop new, creative, and powerful single-molecule analysis methods that fully benefit from probability theory and consistency with the scientific method.

### SUMMARY POINTS

1. In accordance with the scientific method, any analysis of a natural phenomenon requires the application of a model, along with its associated assumptions. The models used to analyze single-molecule biophysics experiments must account for behavior of individual molecules, molecular heterogeneity, and noisy signals.
2. Bayesian inference is the best way to perform the modeling of a single-molecule experiment because it is most consistent with the scientific method and accounts for the uncertainties present in all aspects of the experiments.
3. The use of a prior probability allows the quantitative incorporation of information from previous experiments and theories into the current analysis. It is an integral part of the model and thus should not be dismissed, as is the case in non-Bayesian inference-based methods.
4. Likelihood functions, although integral in relating the model to the observed data, cannot by themselves be used for inference. Doing so addresses the reverse of the problem that the scientific method aims to solve.
5. The posterior probability can be thought of as the ‘updated’ model after performing an experiment. It is the objective of all analysis methods and captures the uncertainty in our knowledge of model parameters.
6. Analyses of single-molecule experiments that use Bayesian model selection (BMS) are able to calculate the probability that a particular model is the ‘best’ model of the underlying natural phenomenon, and therefore allow researchers to quantitatively evaluate hypotheses in a manner that would not otherwise be possible.

### FUTURE ISSUES

1. Wider adoption of existing Bayesian inference-based data analysis methods could greatly benefit the field of single-molecule biophysics. Moreover, wider engagement by the single-molecule biophysics community in extending existing Bayesian inference-based methods and creating new such methods would could be transformative to the field.
2. Many of the complexities of single-molecule behavior and data remain inaccessible to current analysis methods due to the absence of suitable models to describe them. Models capable of describing these behaviors and data, and the experimental techniques used to observe them, need to be developed.
3. A number of currently available Bayesian inference methods are prohibitively expensive in terms of ease of use and/or computational resources required for implementation. More accessible and efficient Bayesian methods need to be developed.
4. Analyses of single-molecule experiments often use Bayesian inference in a piecemeal manner—either for some parts of a larger analysis and/or in a way that has been optimized for a specific type of biomolecule or signal. General, Bayesian inference-based computational frameworks that encompass every part of a single-molecule experiment and are capable of incorporating information from multiple experimental sources to yield a comprehensive picture of the biomolecular process under investigation remain elusive and need to be developed.

### TERMS AND DEFINITIONS

1. **Bayesian model selection (BMS)**: Bayesian inference performed on the evidence of different models to determine the probability that each model is the ‘best’ model.
2. **Evidence**: The probability that the dataset was generated by the given model.
3. **Likelihood function**: The probability that the dataset was generated by particular parameter values according to a given model.
4. **Maximum a posteriori (MAP) estimation**: A Bayesian algorithm where a dataset is modeled using the maximum of the posterior as a point estimate.
5. **Maximum likelihood (ML) estimation**: A model-fitting algorithm where a dataset is modeled using the maximum of the likelihood as a point estimate.
6. **Point estimate**: A location in parameter space used as the best guess of the model parameters (*e.g.*, the maximum of the posterior).
7. **Posterior probability**: The probability that the model parameters can take up particular values upon observation of the data.
8. **Prior probability**: The probability that the model parameters can take up particular values prior to observation of the data.

## ACKNOWLEDGEMENTS

We apologize to all whose work we did not with the illustrative examples chosen here. We thank Chris Wiggins, Jake Hofman, Jonathan Bronson, Jan-Willem van de Meent, and David Blei for many insightful discussions about Bayesian inference over the years, which led to very productive collaborations; Sarah Dubnik, Ethan Feng, and Steve Jang for comments on the manuscript; and Ethan Feng for assistance with the preparation of the illustrations. This work was supported by funds to R.L.G. from the National Institute of Health (R01 GM119386, R01 GM084288, and R01 GM137608), the National Science Foundation (CHE 2004016), a Burroughs Wellcome Fund Career Award in the Biomedical Sciences (1004856), a National Science Foundation CAREER Award (MCB 0644262), an American Cancer Society Research Scholar Award (RSG-09-053-01-GMC), and a Camille Dreyfus Teacher-Scholar Award (DRFSCH CU11-0665). C.D.K-T. acknowledges support from the Department of Energy Office of Science Graduate Fellowship Program (DE-AC05-06OR23100), and Columbia University’s NIH Training Program in Molecular Biophysics (T32-GM008281).

## RELATED RESOURCES

1. Bonomi M, Hanot S, Greenberg CH, Sali A, Nilges M, et al. 2019. Bayesian Weighing of Electron Cryo-Microscopy Data for Integrative Structural Modeling. Structure. 27(1):175–188.e6

2. Dörfler T, Eilert T, Röcker C, Nagy J, Michaelis J. 2017. Structural Information from Single-molecule FRET Experiments Using the Fast Nano-positioning System. JoVE J. Vis. Exp., p. e54782

3. El Beheiry M, Türkcan S, Richly MU, Triller A, Alexandrou A, et al. 2016. A Primer on the Bayesian Approach to High-Density Single-Molecule Trajectories Analysis. Biophys. J. 110(6):1209–15

4. Fredkin DR, Rice JA. 1992. Bayesian Restoration of Single-Channel Patch Clamp Recordings. Biometrics. 48(2):427–48

5. Lindén M, Ćurić V, Amselem E, Elf J. 2017. Pointwise error estimates in localization microscopy. Nat. Commun. 8(1):15115

6. Masson J-B, Casanova D, Türkcan S, Voisinne G, Popoff MR, et al. 2009. Inferring Maps of Forces inside Cell Membrane Microdomains. Phys. Rev. Lett. 102(4):048103

7. Nakane T, Kimanius D, Lindahl E, Scheres SH. 2018. Characterisation of molecular motions in cryo-EM single-particle data by multi-body refinement in RELION. eLife. 7:e36861

8. Rowley MI, Coolen ACC, Vojnovic B, Barber PR. 2016. Robust Bayesian Fluorescence Lifetime Estimation, Decay Model Selection and Instrument Response Determination for Low-Intensity FLIM Imaging. PLOS ONE. 11(6):e0158404

9. Sgouralis I, Pressé S. 2017. An Introduction to Infinite HMMs for Single-Molecule Data Analysis. Biophys. J. 112(10):2021–29

10. Xue XC, Tong H, Liu F, Ou-Yang Z. 2008. Bayesian Analysis of Folding and Unfolding Time Series of Single-Forced RNAs. J. Phys. Chem. B. 112(44):13680–83

## LITERATURE CITED

1. Bishop C. 2006. Pattern Recognition and Machine Learning. New York: Springer-Verlag

2. Bronson JE, Fei J, Hofman JM, Gonzalez RL, Wiggins CH. 2009. Learning Rates and States from Biophysical Time Series: A Bayesian Approach to Model Selection and Single-Molecule FRET Data. Biophys. J. 97(12):3196–3205

3. Chaloner K, Verdinelli I. 1995. Bayesian Experimental Design: A Review. Stat. Sci. 10(3):273–304

4. Cossio P, Hummer G. 2013. Bayesian analysis of individual electron microscopy images: Towards structures of dynamic and heterogeneous biomolecular assemblies. J. Struct. Biol. 184(3):427–37

5. Cox RT. 1946. Probability, Frequency and Reasonable Expectation. Am. J. Phys. 14(1):1–13

6. Du C, Kou SC. 2020. Statistical Methodology in Single-Molecule Experiments. Stat. Sci. 35(1):75–91

7. Ensign DL, Pande VS. 2010. Bayesian Detection of Intensity Changes in Single Molecule and Molecular Dynamics Trajectories. J. Phys. Chem. B. 114(1):280–92

8. Foreman-Mackey D, Hogg DW, Lang D, Goodman J. 2013. emcee: The MCMC Hammer. Publ. Astron. Soc. Pac. 125(925):306–12

9. Goodman J, Weare J. 2010. Ensemble samplers with affine invariance. Commun. Appl. Math. Comput. Sci. 5(1):65–80

10. Habeck M. 2011. Statistical mechanics analysis of sparse data. J. Struct. Biol. 173(3):541–48

11. Habeck M. 2017. Bayesian Modeling of Biomolecular Assemblies with Cryo-EM Maps. Front. Mol. Biosci. 4:

12. Hastings WK. 1970. Monte Carlo sampling methods using Markov chains and their applications. Biometrika. 57(1):97–109

13. Hon J, Gonzalez RL. 2019. Bayesian-Estimated Hierarchical HMMs Enable Robust Analysis of Single-Molecule Kinetic Heterogeneity. Biophys. J. 116(10):1790–1802

14. Jaynes ET. 2003. Probability Theory: The Logic of Science. Cambridge University Press. 762 pp.

15. Johnson S, van de Meent J-W, Phillips R, Wiggins CH, Lindén M. 2014. Multiple LacI-mediated loops revealed by Bayesian statistics and tethered particle motion. Nucleic Acids Res. 42(16):10265–77

16. Karslake JD, Donarski ED, Shelby SA, Demey LM, DiRita VJ, et al. 2020. SMAUG: Analyzing single-molecule tracks with nonparametric Bayesian statistics. Methods, p. S1046202320300293

17. Kimanius D, Zickert G, Nakane T, Adler J, Lunz S, et al. 2020. Exploiting prior knowledge about biological macromolecules in cryo-EM structure determination. bioRxiv. 2020.03.25.007914

18. Kinz-Thompson CD, Gonzalez RL. 2018. Increasing the Time Resolution of Single-Molecule Experiments with Bayesian Inference. Biophys. J. 114(2):289–300

19. Kou SC, Xie XS, Liu JS. 2005. Bayesian analysis of single-molecule experimental data. J. R. Stat. Soc. Ser. C Appl. Stat. 54(3):469–506

20. Kovalevskiy O, Nicholls RA, Long F, Carlon A, Murshudov GN. 2018. Overview of refinement procedures within *REFMAC* 5: utilizing data from different sources. Acta Crystallogr. Sect. Struct. Biol. 74(3):215–27

21. Lartillot N, Philippe H. 2006. Computing Bayes Factors Using Thermodynamic Integration. Syst. Biol. 55(2):195–207

22. Laurent F, Floderer C, Favard C, Muriaux D, Masson J-B, Vestergaard CL. 2019. Mapping spatio-temporal dynamics of single biomolecules in living cells. Phys. Biol. 17(1):015003

23. Lindén M, Elf J. 2018. Variational Algorithms for Analyzing Noisy Multistate Diffusion Trajectories. Biophys. J. 115(2):276–82

24. Malakoff D. 1999. Bayes Offers a “New” Way to Make Sense of Numbers. Science. 286(5444):1460–64

25. Metropolis N, Rosenbluth AW, Rosenbluth MN, Teller AH, Teller E. 1953. Equation of State Calculations by Fast Computing Machines. J. Chem. Phys. 21(6):1087–92

26. Minka TP. 2008. Automatic Choice of Dimensionality for PCA. M.I.T. Media Laboratory Perceptual Computing Section. 514, MIT Media Laboratory, Vision and Modeling Group

27. Monnier N, Barry Z, Park HY, Su K-C, Katz Z, et al. 2015. Inferring transient particle transport dynamics in live cells. Nat. Methods. 12(9):838–40

28. Monnier N, Guo S-M, Mori M, He J, Lénárt P, Bathe M. 2012. Bayesian Approach to MSD-Based Analysis of Particle Motion in Live Cells. Biophys. J. 103(3):616–26

29. Okamoto K, Sako Y. 2012. Variational Bayes Analysis of a Photon-Based Hidden Markov Model for Single-Molecule FRET Trajectories. Biophys. J. 103(6):1315–24

30. Persson F, Lindén M, Unoson C, Elf J. 2013. Extracting intracellular diffusive states and transition rates from single-molecule tracking data. Nat. Methods. 10(3):265–69

31. Robson A, Burrage K, Leake MC. 2013. Inferring diffusion in single live cells at the single-molecule level. Philos. Trans. R. Soc. B Biol. Sci. 368(1611):20120029

32. Rolfe DJ, McLachlan CI, Hirsch M, Needham SR, Tynan CJ, et al. 2011. Automated multidimensional single molecule fluorescence microscopy feature detection and tracking. Eur. Biophys. J. 40(10):1167

33. Scheres SHW. 2012. RELION: Implementation of a Bayesian approach to cryo-EM structure determination. J. Struct. Biol. 180(3):519–30

34. Sgouralis I, Madaan S, Djutanta F, Kha R, Hariadi RF, Pressé S. 2019. A Bayesian Nonparametric Approach to Single Molecule Förster Resonance Energy Transfer. J. Phys. Chem. B. 123(3):675–88

35. Slator PJ, Cairo CW, Burroughs NJ. 2015. Detection of Diffusion Heterogeneity in Single Particle Tracking Trajectories Using a Hidden Markov Model with Measurement Noise Propagation. PLOS ONE. 10(10):e0140759

36. Smith CS, Jouravleva K, Huisman M, Jolly SM, Zamore PD, Grunwald D. 2019. An automated Bayesian pipeline for rapid analysis of single-molecule binding data. Nat. Commun. 10(1):

37. Thapa S, Lomholt MA, Krog J, Cherstvy AG, Metzler R. 2018. Bayesian analysis of single-particle tracking data using the nested-sampling algorithm: maximum-likelihood model selection applied to stochastic-diffusivity data. Phys. Chem. Chem. Phys. 20(46):29018–37

38. Tinoco I, Gonzalez RL. 2011. Biological mechanisms, one molecule at a time. Genes Dev. 25(12):1205–31

39. van de Meent J-W, Bronson JE, Wiggins CH, Gonzalez RL. 2014. Empirical Bayes Methods Enable Advanced Population-Level Analyses of Single-Molecule FRET Experiments. Biophys. J. 106(6):1327–37

40. von Neumann J. 1963. Method in the Physical Sciences. In Collected Works. Volume VI Theory of Games, Astrophysics, Hydrodynamics and Meteorology. VI:491–498. Oxford: Pergamon

41. Zivanov J, Nakane T, Scheres SHW. 2019. A Bayesian approach to beam-induced motion correction in cryo-EM single-particle analysis. IUCrJ. 6(1):5–17

